# Novel mouse model of Weaver syndrome displays overgrowth and excess osteogenesis reversible with KDM6A/6B inhibition

**DOI:** 10.1101/2023.06.23.546270

**Authors:** Christine W Gao, WanYing Lin, Ryan C Riddle, Priyanka Kushwaha, Leandros Boukas, Hans T Björnsson, Kasper D Hansen, Jill A Fahrner

## Abstract

Weaver syndrome is a Mendelian disorder of the epigenetic machinery (MDEM) caused by germline pathogenic variants in *EZH2*, which encodes the predominant H3K27 methyltransferase and key enzymatic component of Polycomb repressive complex 2 (PRC2). Weaver syndrome is characterized by striking overgrowth and advanced bone age, intellectual disability, and distinctive facies. We generated a mouse model for the most common Weaver syndrome missense variant, *EZH2* p.R684C. *Ezh2^R684C/R684C^* mouse embryonic fibroblasts (MEFs) showed global depletion of H3K27me3. *Ezh2^R684C/+^* mice had abnormal bone parameters indicative of skeletal overgrowth, and *Ezh2^R684C/+^* osteoblasts showed increased osteogenic activity. RNA-seq comparing osteoblasts differentiated from *Ezh2^R684C/+^*and *Ezh2^+/+^* bone marrow mesenchymal stem cells (BM-MSCs) indicated collective dysregulation of the BMP pathway and osteoblast differentiation. Inhibition of the opposing H3K27 demethylases Kdm6a/6b substantially reversed the excessive osteogenesis in *Ezh2^R684C/+^* cells both at the transcriptional and phenotypic levels. This supports both the ideas that writers and erasers of histone marks exist in a fine balance to maintain epigenome state, and that epigenetic modulating agents have therapeutic potential for the treatment of MDEMs.

## Introduction

Mendelian disorders of the epigenetic machinery (MDEMs) result from germline variants in writers, erasers, and readers of epigenetic marks, as well as chromatin remodelers (1, 2). The characteristic feature of this class is a combination of growth abnormality (either overgrowth or undergrowth; seen in 74% of these disorders) and developmental delay/intellectual disability (present in 85% of disorders) (3). To date, at least 85 MDEMs have been described, yet treatment still only consists of symptomatic management and preventative screening for known complications (3). Despite being monogenic disorders, MDEMs are thought to cause multisystemic findings through widespread epigenomic dysregulation and consequent transcriptomic perturbation. This has been supported by ATAC-seq and RNA-seq studies in both animal and cell line models of several MDEMs, showing alterations to chromatin accessibility and gene expression (4–6). Recently, peripheral blood DNA methylation signatures have also been identified for many MDEMs, and are now not only approved for diagnosis, but also can reliably differentiate pathogenic variants from benign (7–11). This reinforces the idea that the core etiology of MDEMs lies in their effects upon the epigenome, which is a unifying quality of MDEMs.

Weaver syndrome (MIM 277590) is a MDEM with cardinal signs of overgrowth, developmental delay/intellectual disability, and characteristic facial appearance, as well as advanced osseous maturation (12–16). In 2012, with the advent of whole exome sequencing, Tatton-Brown et al. and Gibson et al. traced the molecular etiology of Weaver syndrome to heterozygous pathogenic variants in *EZH2*, which encodes the primary H3K27 methyltransferase and core component of the Polycomb repressive complex 2 (PRC2) (14, 15). PRC2 is highly conserved across plants, fungi, and animals, and is involved in fundamental processes such as cell differentiation, development, and cell cycle control (17). Variants in other components of PRC2 partially phenocopy Weaver syndrome: Cohen-Gibson syndrome (*EED* variants; MIM 617561) (18–20) and Imagawa-Matsumoto syndrome (*SUZ12* variants; MIM 618786) (21–23) also have overgrowth, advanced bone age, and developmental delay/intellectual disability as central features. Interestingly, all three PRC2 MDEMs share a peripheral blood DNA methylation signature (10), suggesting a common mechanism of disease linked to PRC2 dysfunction.

The exact mechanism by which PRC2 dysfunction translates to an overgrowth phenotype at the organismal level is still being explored. Long bone growth is driven by endochondral ossification at cartilaginous growth plates (24). Chondrocytes at the growth plate undergo rapid proliferation and hypertrophy, during which the cells secrete an abundance of extracellular matrix. This matrix acts as a scaffold for invasion by vasculature, osteoclasts, and osteoblasts. While osteoclasts resorb old matrix, osteoblasts lay down new bone. The fine orchestration of these cell types continues to shape and remodel the bone structure throughout life, and is impacted by changes in lineage commitment, proliferation, and development. This process is also regulated by a plethora of endocrine signals, such as growth hormone (GH) and thyroid hormone, as well as local factors such as Indian hedgehog (IHH), bone morphogenetic proteins (BMPs), and insulin-like growth factors (IGFs), all which in turn influence intracellular transcription regulatory networks. *EZH2* is downregulated within 3 days after the onset of osteogenic differentiation in mesenchymal stem cells (MSCs) (25). Conditional knockout of *Ezh2* in the MSC lineage leads to severe skeletal patterning defects and overexpression of cyclin-dependent kinase (CDK) inhibitors, but siRNA knockdown or pharmacological inhibition of *Ezh2* in MSCs also increases osteogenic marker expression (25, 26). This suggests that *Ezh2* is necessary for early proliferation of osteogenic precursors, yet simultaneously suppresses osteoblast maturation. However, these conditional knockout and complete inhibition models do not capture the growth patterns of Weaver syndrome, which generally originates from constitutive heterozygous pathogenic missense variants.

To investigate the mechanisms behind skeletal overgrowth in Weaver syndrome, we generated a novel constitutive missense mouse model, *Ezh2^R684C/+^*. This heterozygous model both allowed *in vivo* skeletal profiling and provided a source of primary cells for *in vitro* studies of osteoblast differentiation. Here, using micro-computed tomography (micro-CT), we describe a skeletal overgrowth phenotype in *Ezh2^R684C/+^* mice that is reminiscent of Weaver syndrome. Both our *in vivo* labeling and *in vitro* differentiation assays suggest that excessive osteogenesis by the osteoblast lineage bears responsibility for the overgrowth, and we identify a distinct transcriptional profile in *Ezh2^R684C/+^* osteoblasts. Finally, we found reversal of both the osteogenic phenotype and transcriptional perturbations of *Ezh2^R684C/+^* osteoblasts following treatment with an epigenetic modulator, GSK-J4. These findings contribute to the growing body of literature indicating that MDEMs could one day be treated by addressing their epigenetic etiology.

## Results

### A recurrent Weaver syndrome missense variant causes loss of H3K27me3 methyltransferase activity

*EZH2* p.R684C is the most common heterozygous pathogenic variant in unrelated patients with Weaver syndrome (14, 16), and lies within the catalytic SET methyltransferase domain, which is highly conserved across species (27–30). Using CRISPR/Cas9 gene editing, we generated a mouse model bearing the orthologous *Ezh2* missense variant (Figure 1, A and B). We refer to the edited allele as *Ezh2^R684C^* hereafter for simplicity. *Ezh2^R684C/+^* pups were born full-term at Mendelian ratios (Supplemental Table 1, A–C). This suggests that a drastic disruption to global H3K27me3 is unlikely in the heterozygous state, since H3K27me3 is a critical mark for transcriptional regulation, maintenance of facultative heterochromatin, and thus initiation and preservation of cell differentiation state (31). Embryos homozygous for the R684C variant (*Ezh2^R684C/R684C^*) were viable through E14.5, but ultimately did not survive to birth (Supplemental Table 1, A and B).

**Figure 1:**
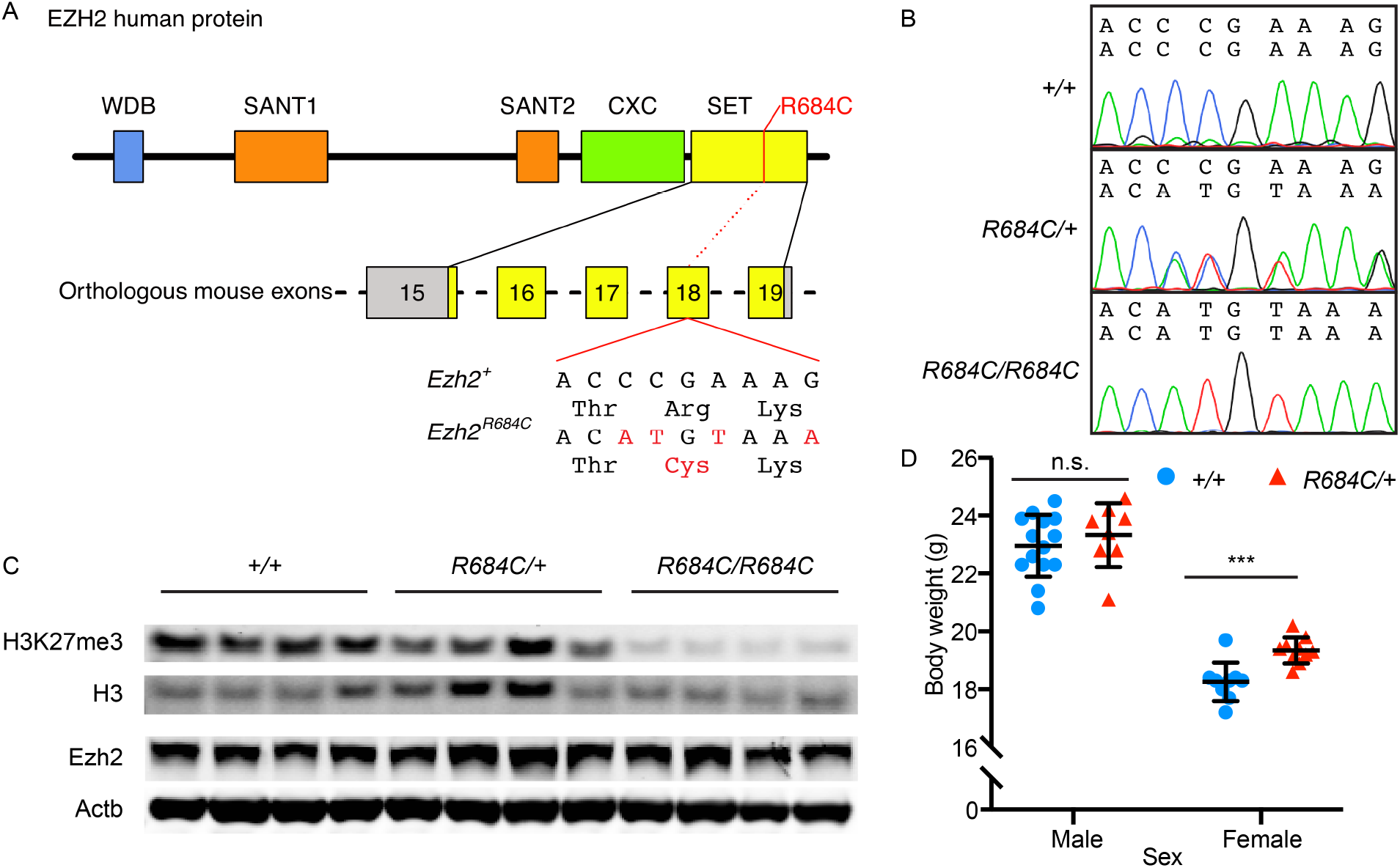
The *Ezh2* R684C allele leads to reduced H3K27me3 catalysis and overgrowth in female mice. (A) Four base changes were introduced into exon 18 of *M. musculus Ezh2*, changing codon CGA (Arg 679) in the catalytic SET domain (yellow) to TGT (Cys) and introducing two silent mutations to create an Nsp1 restriction site for genotyping. At the protein level, this corresponds to *H. sapiens* EZH2 p.R684C. (B) Chromatogram traces for E14.5 mouse embryonic fibroblasts (MEFs) that are wild-type at the *Ezh2* locus (+/+), heterozygous (*R684C/+*), or homozygous for the *R684C* variant allele (*R684C/R684C*). (C) Western blot showing that *R684C/R684C* MEFs have reduced global H3K27me3, which is minimally changed in *R684C/+* MEFs compared to *+/+* MEFs. All three genotypes have comparable levels of Ezh2. H3 and Actb served as loading controls. (D) Female *Ezh2^R684C/+^* mice have increased body weight at 8 weeks of age compared to female *Ezh2^+/+^* littermates. *Ezh2^+/+^* males n=14, *Ezh2^+/+^* females n=9. *Ezh2^R684C/+^* males n=8, *Ezh2^R684C/+^* females n=10. Blue circles represent *Ezh2^+/+^*; red triangles represent *Ezh2^R684C/+^*. ***p < 0.001, unpaired Student’s t-test. All error bars represent mean ±1 SD. n.s., non-significant.

*Ezh2^R684C/R684C^* mouse embryonic fibroblasts (MEFs) isolated at E14.5 had wild-type levels of Ezh2 protein expression (Figure 1C). This indicated that the R684C allele produces a full-length protein product, albeit catalytically compromised, as H3K27me3 levels were drastically decreased (Figure 1C). Minimal residual H3K27me3 in *Ezh2^R684C/R684C^* MEFs may be due to Ezh1, a homolog of Ezh2 that has low levels of H3K27me3 methyltransferase activity (32). Our data show that R684C is a hypomorphic variant, or possibly loss-of-function, that interferes with SET domain catalysis but does not affect protein stability. Interestingly – and consistent with the aforementioned lack of alteration of Mendelian birth ratios – global H3K27me3 levels were minimally altered on western blot in heterozygous *Ezh2^R684C/+^* MEFs. This genotype therefore appears able to sustain global H3K27 tri-methyltransferase activity at near-normal levels. It is thus likely that subtle shifts in H3K27me3 levels at particular genomic loci, with ensuing shifts in gene expression, may play a role in the phenotype of this mouse model. Such local alterations may be undetectable by western blot.

### *Ezh2^R684C/+^* mice exhibit a skeletal overgrowth phenotype

Similar to the Weaver syndrome phenotype, *Ezh2^R684C/+^* mice displayed overgrowth. Female *Ezh2^R684C/+^* mice showed significantly higher body weight compared to *Ezh2^+/+^* littermates at 8 weeks of age (p=0.0006), although we did not see growth differences in male *Ezh2^R684C/+^* mice (Figure 1D and Supplemental Figure 1). Direct measurements and high-resolution micro-computed tomography (micro-CT) imaging did not reveal a difference in femur and tibia length between 8 week-old *Ezh2^R684C/+^* and *Ezh2^+/+^* mice (Figure 2, A and B, and Supplemental Figure 2A), nor did we observe differences in total body length (Supplemental Figure 2B), or trabecular bone structure in the distal femur (Supplemental Figure 2, C–E). However, *Ezh2^R684C/+^* mice did exhibit striking alterations in cortical bone structure at the femoral mid-diaphysis. Cross-sectional tissue area was significantly increased in both male and female *Ezh2^R684C/+^* mice when compared to *Ezh2^+/+^* littermates (p=0.032 and p<1×10^−6^, respectively) (Figure 2, C and D), which led to a decrease in the bone area/tissue area percentage in females (p=0.028) (Figure 2E). Female *Ezh2^R684C/+^* mice also had a trend towards increased cortical thickness (p=0.057) (Figure 2F).

**Figure 2:**
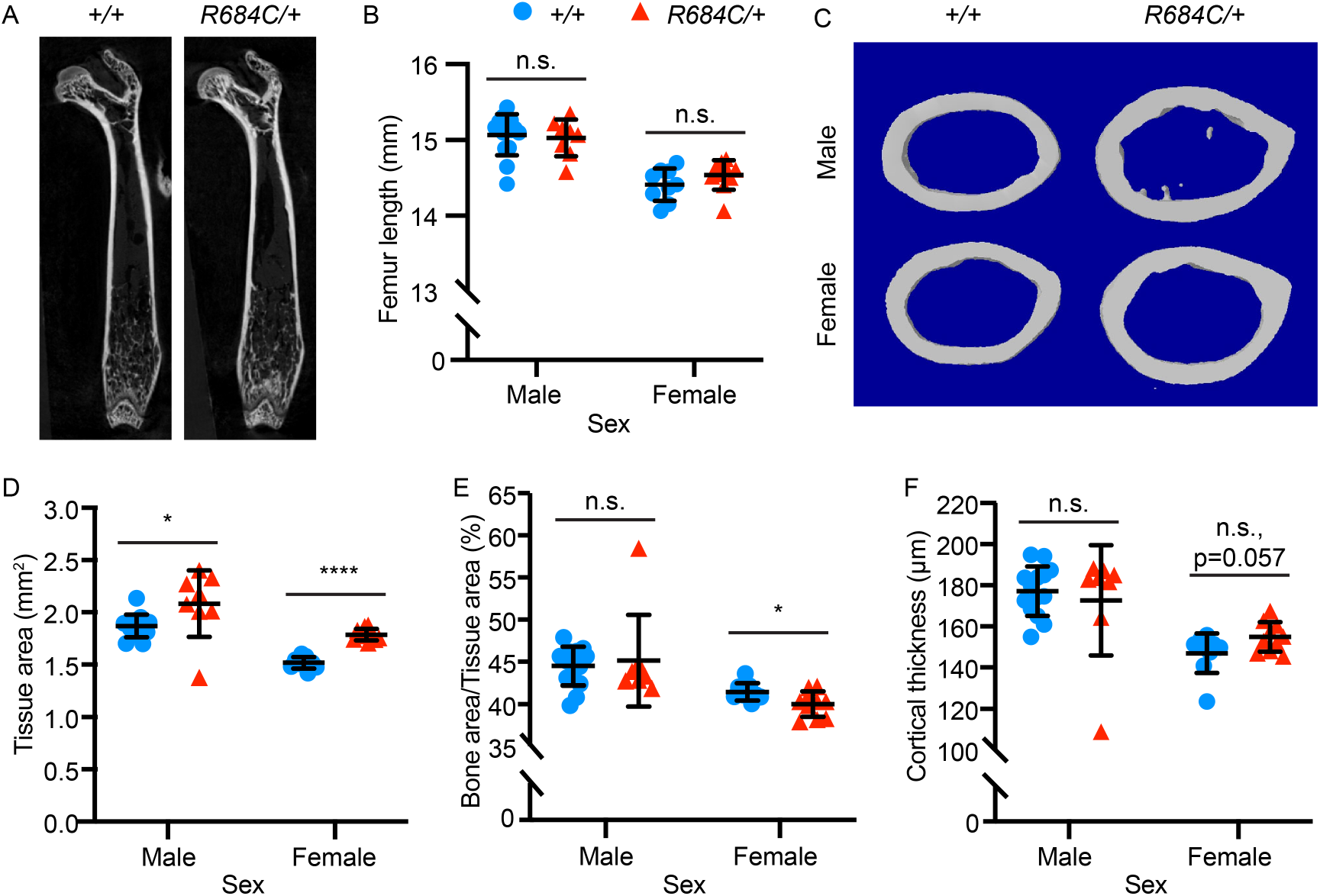
*Ezh2^R684C/+^* mice have altered cortical bone parameters. (A) Micro-computed tomography (micro-CT) of *Ezh2^+/+^* and *Ezh2^R684C/+^* femurs in the coronal plane. (B) Femur lengths do not differ between *Ezh2^+/+^* and *Ezh2^R684C/+^* mice for either sex. (C) Micro-CT reconstructions of cortical bone regions of interest at the mid-diaphysis. (D) Tissue area is significantly increased in *Ezh2^R684C/+^* mice of both sexes. (E) *Ezh2^R684C/+^* female mice have a lower bone area/tissue area percentage. (F) Female *Ezh2^R684C/+^* mice have a trend towards higher cortical thickness (n.s., p=0.057). *Ezh2^+/+^* males n=14, *Ezh2^+/+^* females n=9. *Ezh2^R684C/+^* males n=8, *Ezh2^R684C/+^* females n=10. Blue circles represent *Ezh2^+/+^*; red triangles represent *Ezh2^R684C/+^*. *p < 0.05, ****p < 0.0001, unpaired Student’s t-test. All error bars represent mean ±1 SD. n.s., non-significant.

To understand the basis for the cortical bone phenotype, we quantified the rate of bone formation by dynamic histomorphometry after sequential injection of calcein and Alizarin Red at 5 weeks of age, when bone growth is rapid, yet prior to discernable overgrowth. The mineral apposition rate (MAR) is a direct measure of osteoblast activity. Compared to *Ezh2^+/+^* littermates, periosteal MAR was significantly increased in female *Ezh2^R684C/+^* mice (p=0.001) (Figure 3, A and B), and endosteal MAR was increased in both male and female *Ezh2^R684C/+^* mice (p=0.033 and p=0.028, respectively) (Figure 3, A and C). These findings indicate that the *Ezh2^R684C/+^* mouse model recapitulates certain overgrowth aspects of Weaver syndrome, and further suggest that cortical bone modeling is altered in *Ezh2^R684C/+^* mice via excessive osteoblast activity.

**Figure 3:**
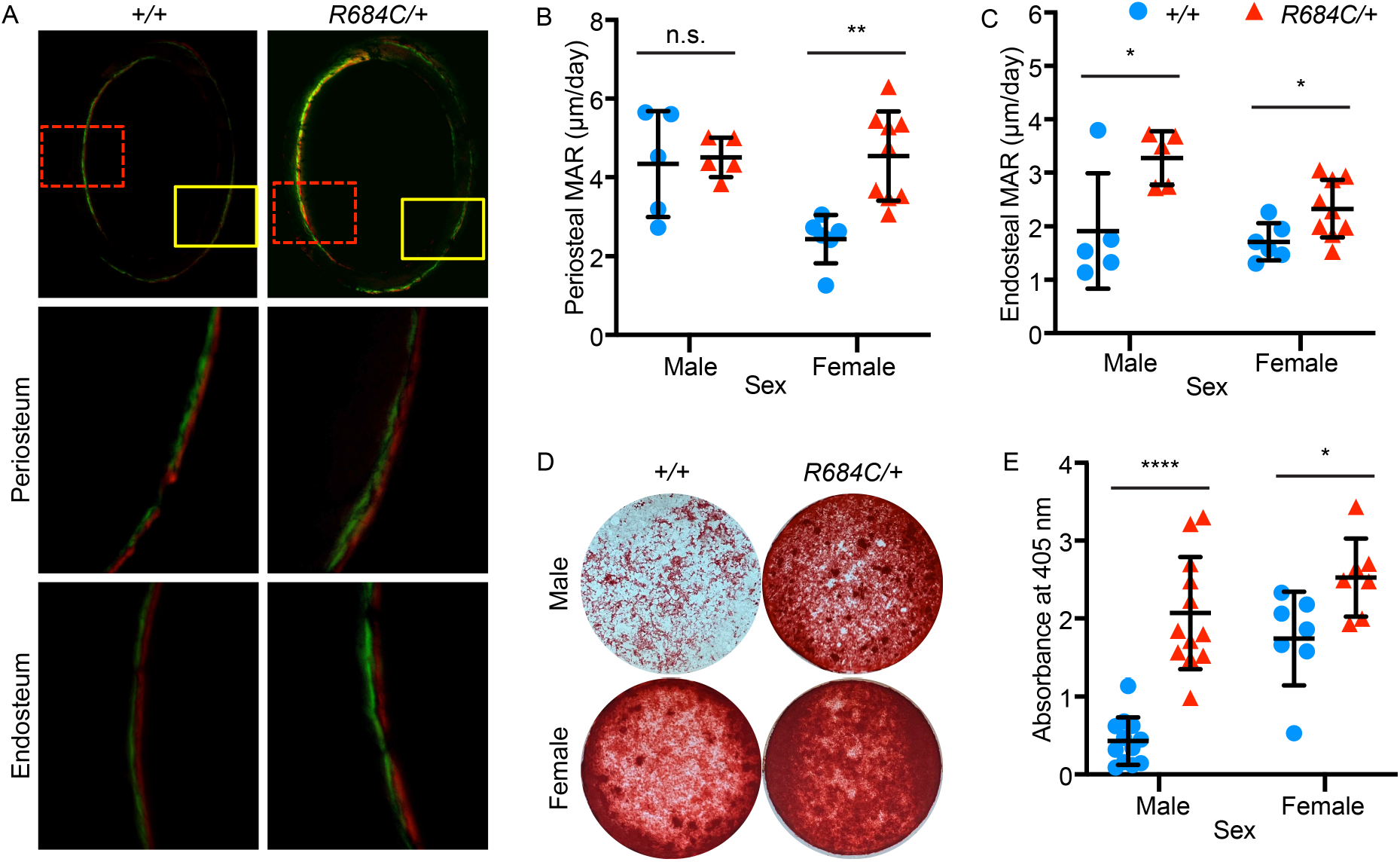
Osteoblast activity is increased in *Ezh2^R684C/+^* mice. (A) Representative images of double-fluorescent *in vivo* labeling at the femoral mid-diaphysis, in the transverse plane (4x magnification, top panels). Green – calcein; red – Alizarin red S. Solid yellow boxes mark the regions for periosteal measurements (20x magnification, center panels); dashed red boxes mark the regions for endosteal measurements (20x magnification, bottom panels). Mineral appositional rate (MAR) is increased at the (B) periosteum in females only, and at the (C) endosteum for both sexes. *Ezh2^+/+^* males n=5, *Ezh2^+/+^* females n=6. *Ezh2^R684C/+^* males n=5, *Ezh2^R684C/+^* females n=9. (D) Alizarin red staining of osteoblasts following 21 days of *in vitro* differentiation from primary murine bone marrow mesenchymal stem cells (BM-MSCs) isolated from *Ezh2^R684C/+^* and *Ezh2^+/+^* mice. Representative whole-well images taken from a 24-well plate. (E) *Ezh2^R684C/+^* cells of either sex have higher uptake of Alizarin red, as quantified by absorbance at 405 nm. *Ezh2^+/+^* males n=12, *Ezh2^+/+^* females n=7, *Ezh2^R684C/+^* males n=12, *Ezh2^R684C/+^*females n=7. Blue circles represent *Ezh2^+/+^*; red triangles represent *Ezh2^R684C/+^*. *p < 0.05, **p < 0.01, ****p < 0.0001, unpaired Student’s t-test. All error bars represent mean ±1 SD. n.s., non-significant.

### *Ezh2^R684C/+^* bone marrow mesenchymal stem cells exhibit higher osteogenic potential *in vitro*

To further investigate the pathogenic role of *Ezh2^R684C/+^* osteoblasts *in vitro*, we isolated murine bone marrow mesenchymal stem cells (BM-MSCs) from 8-10 week-old mice and differentiated the cultures to osteoblasts (33). After 21 days of differentiation in identical conditions, *Ezh2^R684C/+^* cultures had significantly increased uptake of Alizarin red compared to *Ezh2^+/+^* in both males and females, indicative of enhanced deposition of calcium and mineralized bone matrix (males: p<1×10^−6^, females: p=0.021) (Figure 3, D and E). This occurred in the setting of stable cell numbers (data not shown). qPCR confirmed increased expression of osteogenic markers in *Ezh2^R684C/+^* osteoblasts, and even in the undifferentiated BM-MSC state for certain genes (Supplemental Figure 3, A–H), suggesting that *Ezh2^R684C/+^* BM-MSCs may be primed towards osteogenic differentiation. Altogether, these results support our *in vivo* findings and indicate that osteoblasts are critical to the *Ezh2^R684C/+^* phenotype, and also demonstrated the use of BM-MSCs as an *in vitro* model of osteoblast differentiation for further studies.

### Osteoblasts differentiated from *Ezh2^R684C/+^* BM-MSCs have a distinct gene expression profile

To identify gene expression changes responsible for enhanced osteogenesis in Weaver syndrome, we differentiated *Ezh2^R684C/+^* and *Ezh2^+/+^* BM-MSCs towards osteoblasts for 14 days, and then performed transcriptome profiling with RNA-seq. We chose this time point because transcriptional changes are usually evident prior to the appearance of phenotypic differences such as enhanced osteogenesis. Six female biological replicates were sequenced per genotype; principal component analysis indicated that samples clustered approximately according to genotype (Figure 4A). A histogram of gene-wise p-values displayed a non-uniform distribution, with an overrepresentation of low p-values (Figure 4B). As a lack of differentially expressed genes would have resulted in a uniform distribution, this indicated the presence of differential gene expression in *Ezh2^R684C/+^* osteoblasts. In total, 194 genes were differentially expressed at the 10% FDR level (Supplemental Appendix 1), comprising of 94 upregulated genes and 100 downregulated genes, referenced against *Ezh2^+/+^* osteoblasts (Supplemental Figure 4A). Of these, 36 genes had an absolute fold-change greater than 2, indicating a major difference in expression between *Ezh2^R684C/+^* and *Ezh2^+/+^*. We did not see a systemic up-regulation of gene expression in *Ezh2^R684C/+^* cells, which corroborates our finding that H3K27me3 levels do not exhibit major global alteration. We verified that the differentially expressed genes we found in mouse osteoblasts roughly correspond to known human EZH2 target genes (Supplemental Figure 4B). Some incongruence is present, as expected due to cell type and species differences. We also cross-validated our data with a publicly available RNA-seq dataset that investigated loss of Ezh2 function through pharmacological inhibition with GSK-126 (34). This showed that genes affected by Ezh2 inhibition are enriched among the population of genes perturbed in *Ezh2^R684C/+^* cells, and vice versa (Supplemental Figure 4, C and D).

**Figure 4:**
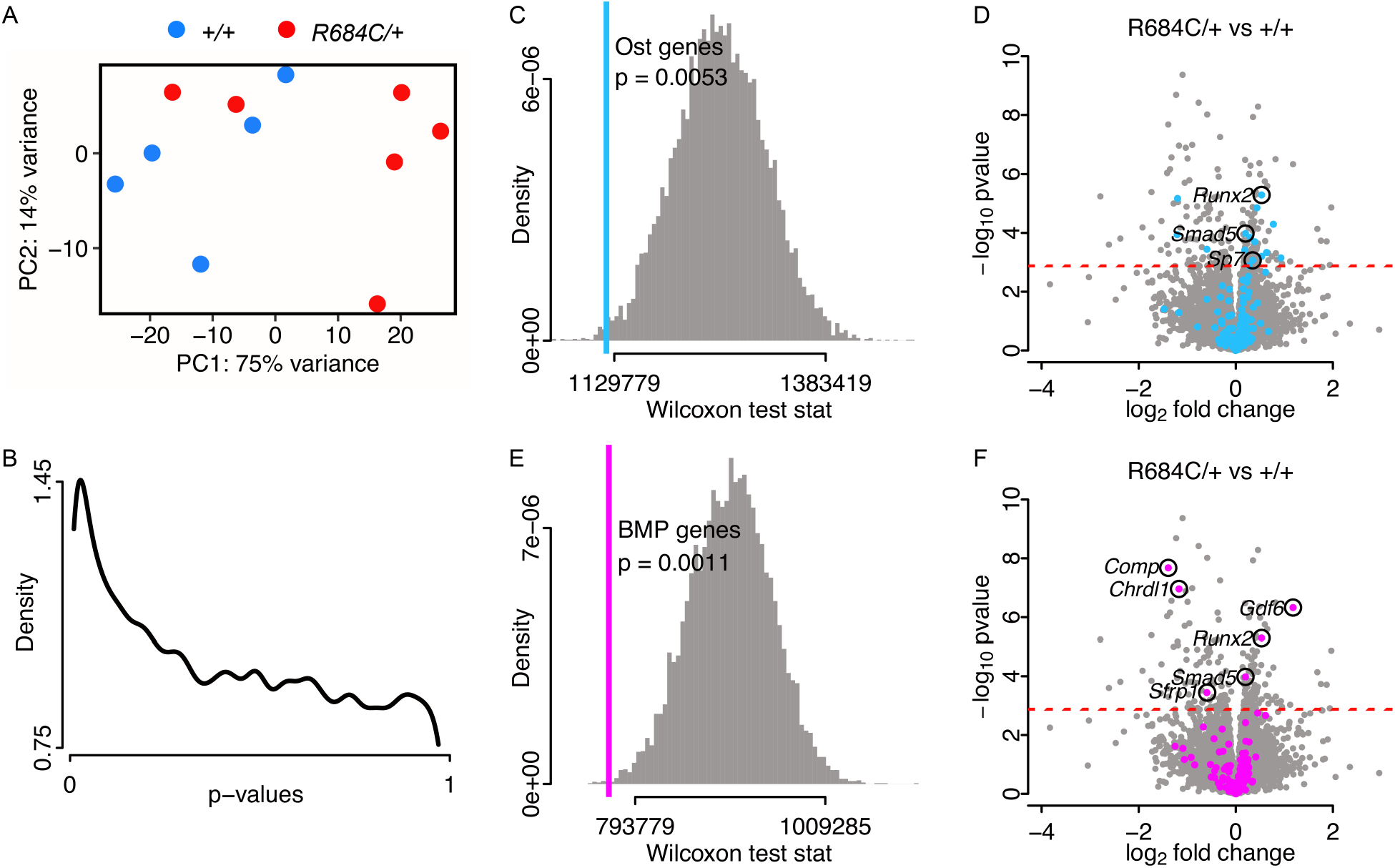
*Ezh2^R684C/+^* osteoblasts demonstrate transcriptional dysregulation of key osteogenic pathways. (A) Principal component analysis of *Ezh2^+/+^* (blue circles, n=5) and *Ezh2^R684C/+^* (red circles, n=6) RNA-seq samples at day 14 of osteoblast differentiation. (B) Density plot of p-values from differential expression analysis comparing *Ezh2^R684C/+^* versus *Ezh2^+/+^* samples, indicating an enrichment of low p-values. (C) Wilcoxon test statistic for Mouse Genome Informatics (MGI) osteoblast differentiation genes (blue line, p=0.0053) plotted over the simulated test statistic distribution for 10,000 random groupings of genes. (D) Volcano plot for *Ezh2^R684C/+^* versus *Ezh2^+/+^* samples, with MGI osteoblast differentiation genes highlighted in blue. False discovery rate (FDR) = 0.1 (red dashed line). (E) Wilcoxon test statistic and simulated test statistic distribution for MGI BMP pathway genes (magenta, p=0.0011). (F) Volcano plot comparing *Ezh2^R684C/+^* vs *Ezh2^+/+^* samples. MGI BMP pathway genes highlighted in magenta. FDR = 0.1 (red dashed line). Ost., osteoblast.

Because osteoblast differentiation is proximal to the mineralization phenotype observed upon *in vitro* Alizarin red staining as well as our prior *in vivo* dynamic histomorphometry data, we focused on this cell type. We utilized Gene Ontology (GO) annotations maintained by Mouse Genome Informatics (MGI) to compile a list of 201 genes involved in osteoblast differentiation (Supplemental Appendix 4), of which 179 genes were expressed in our dataset. The p-value ranks of these selected genes were significantly shifted from the p-value rank of 179 genes randomly selected from our dataset (p=0.0053) (Figure 4C). This suggests that differentiation was collectively dysregulated in *Ezh2^R684C/+^* osteoblasts, even though not every individual gene had detectable dysregulation. Several key pro-osteogenic transcriptional factors were upregulated, such as *Runx2*, *Smad5*, and *Sp7*, and these genes warrant further study (Figure 4D). When we correlated the counts per million (CPM) for each of the osteoblast differentiation genes with the first principal component of the gene expression matrix (PC1), we found high correlation coefficients for this subset of genes compared to selecting random groups of genes (Supplemental Figure 5). This suggests that genes involved with osteoblast differentiation are a key contributor towards the variation captured by PC1 and therefore the distinction between *Ezh2^R684C/+^* and *Ezh2^+/+^* cells.

The BMP pathway is known to play an important role in osteogenesis, and we noticed an apparent abundance of associated genes among RNA-seq hits. Therefore, we performed a similar analysis using an MGI-curated list of 161 BMP pathway genes (Supplemental Appendix 5) and discovered that this process is likewise dysregulated (p=0.0011) (Figure 4E). *Gdf6*, *Runx2*, *Smad5*, *Comp*, *Chrdl1*, and *Sfrp1* are of particular interest for future investigation, as they are differentially expressed (Figure 4F). Notably, 38 genes are annotated as part of both the BMP pathway and osteoblast differentiation, as the processes are intricately intertwined. These pathway analyses provide further support, at a transcriptional level, of perturbed osteogenesis in *Ezh2^R684C/+^* osteoblasts. Overall, our RNAseq results also paint a profile of numerous shifts in transcription acting in concert to produce a phenotype.

### Inhibition of H3K27me3 demethylases with GSK-J4 corrects excessive osteogenesis *in vitro*

Because Ezh2 is the primary methyltransferase (writer) of H3K27me3, we reasoned that the characteristic features of Weaver syndrome might be attributed to decreased or displaced H3K27me3 at key genomic loci. A potential therapeutic strategy, therefore, could involve re-balancing the epigenome through inhibition of the opposing H3K27me3 demethylases (erasers) (Figure 5A). We treated female *Ezh2^R684C/+^* and *Ezh2^+/+^* BM-MSCs with either DMSO (vehicle) or GSK-J4, which is a dual inhibitor of Kdm6a and Kdm6b (35), the primary H3K27me3 erasers in mice and humans (36, 37). Vehicle-treated *Ezh2^R684C/+^* cells had significantly increased Alizarin red staining after 21 days of osteoblast differentiation compared to vehicle-treated *Ezh2^+/+^* cells (p-adj=0.022) (Figure 5, B and C), reinforcing our earlier finding in untreated osteoblasts (Figure 3, D and E). For both *Ezh2^R684C/+^* and *Ezh2^+/+^*, cells treated with 2 µM of GSK-J4 during Days 0-7 of differentiation had significantly decreased Alizarin red staining compared to vehicle-treated cells of the same genotype (p-adj=0.030 and 0.030 for *Ezh2^+/+^* and *Ezh2^R684C/+^*, respectively) (Figure 5, B and C). There was no observed decrease in cell viability due to GSK-J4 treatment at 2 µM (Supplemental Figure 6, A and B). Importantly, GSK-J4-treated *Ezh2^R684C/+^* cells showed no statistically significant difference in Alizarin red staining compared to DMSO-treated *Ezh2^+/+^* cells (p-adj=0.598). This suggests that GSK-J4 can reverse the excessive osteogenesis of *Ezh2^R684C/+^* osteoblasts towards the wild-type baseline. Similar results were found in male *Ezh2^R684C/+^* and *Ezh2^+/+^* osteoblasts (Supplemental Figure 6, C and D).

**Figure 5:**
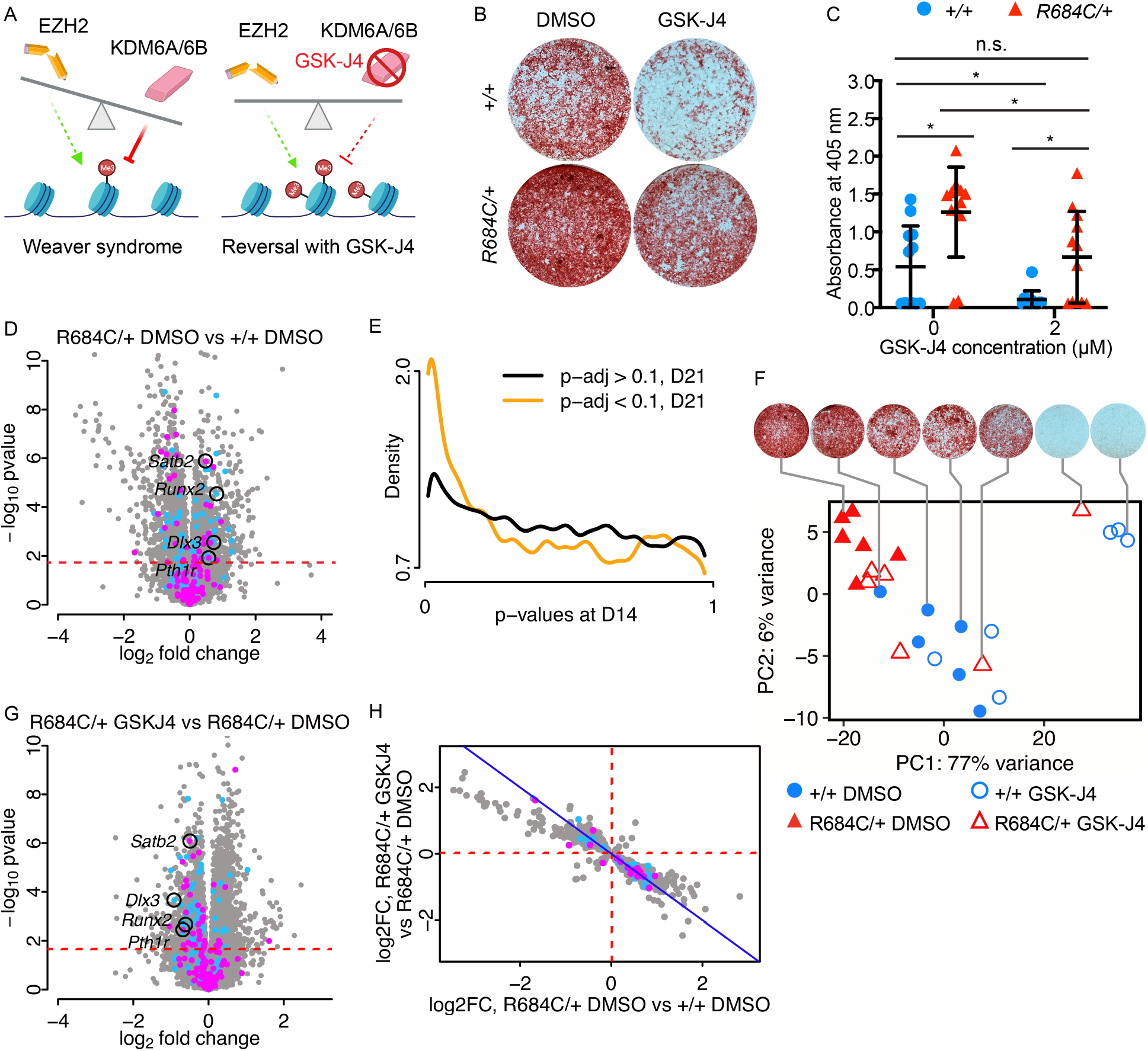
The Kdm6a/6b inhibitor GSK-J4 substantially reverses the *Ezh2^R684C/+^* osteogenic phenotype and transcriptomic profile. (A) Balance hypothesis (1). Left: loss of EZH2 in Weaver syndrome allows for unopposed demethylase activity by KDM6A/6B. Right: inhibition of KDM6A/6B by GSK-J4 restores balance to the chromatin state. (B) Alizarin red staining of female *Ezh2^R684C/+^* and *Ezh2^+/+^* osteoblasts treated with 2 µM GSK-J4 or vehicle (DMSO). Cells were differentiated for 21 days from BM-MSCs. Representative whole-well images shown. (C) GSK-J4 treatment decreases Alizarin red staining in *Ezh2^R684C/+^* and *Ezh2^+/+^* osteoblasts, as quantified by absorbance at 405 nm. *Ezh2^R684C/+^*osteoblasts continue to have higher absorbance than *Ezh2^+/+^*. No significant difference between *Ezh2^+/+^*vehicle-treated and *Ezh2^R684C/+^* GSK-J4-treated osteoblasts. Blue circles: *Ezh2^+/+^* females n=12. Red triangles: *Ezh2^R684C/+^* females n=12. *p < 0.05, unpaired Student’s t-test with Benjamini-Hochberg procedure. All error bars represent mean ±1 SD. n.s., non-significant. (D) Volcano plot of day 21 RNA-seq, displaying log_2_ fold-changes in the *Ezh2^R684C/+^* DMSO vs *Ezh2^+/+^* DMSO contrast. Blue: MGI osteoblast differentiation genes. Magenta: MGI BMP pathway genes. FDR = 0.1 (red dashed line). (E) Conditional p-value density plot displaying p-values from the day 14 untreated RNA-seq, stratified by significance at day 21 in the *Ezh2^R684C/+^* DMSO vs *Ezh2^+/+^* DMSO contrast (orange line, p-adj < 0.1) or not (black line, p-adj > 0.1). (F) Principal component analysis of *Ezh2^+/+^* (blue circles, n=5) and *Ezh2^R684C/+^* (red triangles, n=6) RNA-seq samples at day 21 of osteoblast differentiation, treated either with vehicle (filled icons) or GSK-J4 (open icons). Corresponding Alizarin red staining images shown for representative samples. (G) Volcano plot of day 21 RNA-seq, displaying log_2_ fold-changes in the *Ezh2^R684C/+^* GSKJ4 vs *Ezh2^R684C/+^* DMSO contrast. FDR=0.1 (red dashed line). (H) Scatter plot comparing log_2_ fold-changes in the *Ezh2^R684C/+^* DMSO vs *Ezh2^+/+^* DMSO contrast and corresponding log_2_ fold-changes in the *Ezh2^R684C/+^* GSKJ4 vs *Ezh2^R684C/+^* DMSO contrast. Only genes meeting an adjusted p-value threshold corresponding to FDR < 0.1 in both contrasts are shown (n=1,075). Blue: MGI osteoblast differentiation genes. Magenta: MGI BMP pathway genes.

### GSK-J4 alters the transcriptional profile of *in vitro* differentiated *Ezh2^R684C/+^* osteoblasts

To determine the molecular basis of the GSK-J4 therapeutic effect, we investigated whether GSK-J4 could reverse the transcriptional state of *Ezh2^R684C/+^* cells towards a wild-type profile. To this end, we performed RNA sequencing on *Ezh2^R684C/+^* and *Ezh2^+/+^* osteoblasts. Cells from 6 female mice per genotype were treated with either DMSO (vehicle) or 2 µM of GSK-J4 for the first 7 days of differentiation, and harvested on day 21. Comparison of vehicle-treated *Ezh2^R684C/+^* (*R684C/+* DMSO) to vehicle-treated *Ezh2^+/+^* (*+/+* DMSO) osteoblasts yielded 2,659 differentially expressed genes at the 10% FDR level (Figure 5D) (Supplemental Appendix 2). This is more than was observed between genotypes in our earlier RNA-seq on day 14 untreated cells (Supplemental Figure 4A), and is not accounted for by differences in mean read counts per gene (Supplemental Figure 7A) or coefficients of variation (Supplemental Figure 7, B–D). Reassuringly, among the abundance of significant hits in the *R684C/+* DMSO versus *+/+* DMSO comparison, there was a clear enrichment of genes previously identified as differentially expressed at day 14 (Figure 5E). Based on these results, we surmise that between day 14 and day 21, when the final osteoblast differentiation state is achieved, genotype-specific effects create a greater differential in transcriptional profile between *Ezh2^R684C/+^* and *Ezh2^+/+^* cells.

Principal component analysis of all samples showed that PC1 accounted for 77% of gene expression variance and was closely correlated with corresponding Alizarin red quantifications for the same cell line and treatment condition (r = −0.89) (Figure 5F). To verify that the RNA-seq data was showing an osteogenic transcriptional signal, we also ran linear correlations between Alizarin red quantifications and individual gene counts per million. Among the 20 genes with the highest positive correlation coefficients were *Col1a1*, *Col1a2*, *Smad3*, and *Cthrc1*, indicating that the RNA-seq captured an osteogenic transcriptional profile in samples with high uptake of Alizarin red (38–40).

The *R684C/+* DMSO versus *+/+* DMSO contrast represents genes that are perturbed by the R684C allele. An ideal therapeutic would reverse the transcriptional effects of R684C while minimizing disturbance to the rest of the transcriptome, although this efficacy and specificity is difficult to achieve in practice. The effect of GSK-J4 upon the *Ezh2^R684C/+^* phenotype is captured by comparing GSK-J4-treated *Ezh2^R684C/+^* cells (*R684C/+* GSKJ4) to vehicle-treated *Ezh2^R684C/+^* cells (*R684C/+* DMSO). This contrast yielded an upregulation of 1,562 genes in the *R684C/+* GSKJ4 condition and a downregulation of 1,484 genes, for a total of 3,046 differentially expressed genes at the 10% FDR level (Figure 5G) (Supplemental Appendix 3). Of these, 1,075 genes were also differentially expressed in the *R684C/+* DMSO vs *+/+* DMSO contrast, i.e. both perturbed by the R684C allele and targeted by GSK-J4. Most remarkably, 1,045 of the 1,075 shared genes appear to reverse their fold-change directionality with GSK-J4 treatment (Figure 5H), including 19 genes in the BMP pathway and 31 genes involved in osteoblast differentiation (8 of these overlap). For example, *Satb2*, *Runx2*, *Dlx3*, and *Pthr1* are upregulated in the *R684C/+* DMSO condition relative to *+/+* DMSO, but become downregulated upon GSK-J4 treatment (Figure 5, D and G). These results indicate that GSK-J4 is able to reverse much of the altered transcriptional profile of *Ezh2^R684C/+^* cells towards a wild-type state.

## Discussion

Here we developed and characterized a novel mouse model for the most commonly encountered pathogenic variant in Weaver syndrome, *EZH2* p.R684C. Individuals with Weaver syndrome are often tall, with heights >2 SDs above the age-matched mean, and have additional skeletal abnormalities such as advanced osseous maturation and metaphyseal widening (13, 14, 16). *Ezh2^R684C/+^* mice display skeletal overgrowth and bone structural abnormalities, which are reminiscent of Weaver syndrome. We show this overgrowth to be driven by hyperactivity of the osteoblast lineage in mice. In addition, BM-MSCs isolated from *Ezh2^R684C/+^* mice undergo excessive osteogenesis upon and even prior to differentiation, which was reflected both at the levels of the transcriptome and the cellular phenotype. This may underlie the skeletal overgrowth and advanced bone age observed in many individuals with Weaver syndrome.

### Overgrowth is more subtle in mouse models of Weaver syndrome than in human individuals

While extreme tall stature is a frequent manifestation of Weaver syndrome, with some individuals achieving heights up to +8.3 SD from the age-matched population mean (41), this feature is less prominent in *Ezh2^R684C/+^* mice. We did not observe a significant difference in femur, tibia, or total body length compared to *Ezh2^+/+^* littermates, and only female *Ezh2^R684C/+^* mice had increased weight at 8 weeks. Our results are consistent with those observed in a mouse model for another Weaver syndrome missense variant, V626M (42). In these *Ezh2^V626M/+^* mice, tibia length was also unchanged, and there was a sexually dimorphic effect on weight, with female *Ezh2^V626M/+^* mice having a greater increase in weight. Overgrowth has not been reported to be more pronounced in female individuals with Weaver syndrome, although this may be due to a lack of systematic investigation, as reported cases of Weaver syndrome number <100.

The majority of MDEMs have growth abnormalities, and epigenetic machinery (EM) genes are overrepresented as a cause of overgrowth (43). This implies that genes involved in skeletal growth may be under a greater degree of control by epigenetic mechanisms, and thus particularly vulnerable to imbalances in the epigenetic machinery. Yet, it has been challenging to model overgrowth of MDEMs in mice. The first heterozygous *Ezh2* knockout mouse model appeared to have no phenotype, although in-depth skeletal profiling was not performed (44). Sotos syndrome (MIM 117550) is by far the most common MDEM with overgrowth, caused by variants in *NSD1* as well as 5q35 microdeletions that span *NSD1* (43). However, mice heterozygous for the syntenic microdeletion were found to have decreased weight *in utero* and at 28 weeks of age (45). *Nsd1^+/−^* mice, with loss of one allele due to a premature stop codon, display some characteristic behavioral abnormalities but do not have overt differences in body weight (46). More detailed profiling with micro-CT may nevertheless shed light on features of skeletal overgrowth, as in our *Ezh2^R684C/+^* mice. Transgenic mice are invaluable tools in elucidating the mechanisms of rare disorders; however, it is also important to remain cognizant of potential differences in skeletal growth regulation between mice and humans.

### R684C appears to be a loss-of-function/hypomorphic variant

The mechanism by which *EZH2* variants cause Weaver syndrome has been a matter of discussion. Almost all patients bear heterozygous missense variants (14–16, 41). Cohen et al. previously showed *in vitro* that EZH2 p.R684C, as well as several other missense variants reported in Weaver syndrome, had reduced incorporation of ^3^H-S-adenosyl-methionine (^3^H-SAM) onto core histones (47). We showed that neither *Ezh2^R684C/+^* nor *Ezh2^R684C/R684C^* MEFs have decreased Ezh2 protein levels, yet we observed that *Ezh2^R684C/R684C^* MEFs experience a drastic loss of H3K27me3. There was no increase in H3K27me3 in *Ezh2^R684C/+^* MEFs to suggest paradoxical hypermorph activity, which is known to occur with certain somatic *EZH2* variants found in B-cell lymphomas (48, 49). Therefore, we conclude that R684C is a catalytic loss-of-function/hypomorphic variant (with trace H3K27me3 in the homozygous state possibly due to remaining Ezh1 activity).

In contrast, a recent study instead suggested that R684C could have a dominant negative mechanism. Deevy et al. performed lentiviral transduction of *Ezh2* p.R684C, as well as other Weaver syndrome missense variants, into mouse embryonic stem cells (ESCs) that were heterozygous knockout for *Ezh2* (*Ezh2^fl/Δ^*) (50). The global decrease in H3K27me3 resulting from the missense variants was notably more severe than in non-transduced *Ezh2^fl/Δ^*, which was corroborated by H3K27me3 ChIP-seq. Discrepancies between our findings and those of Deevy et al. on the R684C variant may be due to differences in our model systems, as we used primary MEFs isolated from *Ezh2^R684C/+^* mice at a considerably later developmental stage than ESCs. In the future, it would be interesting to investigate loci-specific H3K27me3 occupancy as well in our system, and especially in osteoblasts.

### Haploinsufficient variants are underrepresented in Weaver syndrome

While most MDEMs result from haploinsufficiency (2), the extreme preponderance of missense variants in Weaver syndrome over frameshift, nonsense, and whole-gene deletion variants suggests an alternate mechanism. One possibility is that manifestation of the Weaver phenotype may depend upon maintaining normal levels of EZH2 protein production, even if reduced in catalytic capacity. Proteins frequently bear multiple functions, known as “moonlighting” (51). EZH2 is known for its H3K27 methyltransferase activity, but it may also play a functional role independent of catalysis. Only 4 *EZH2* variants leading to premature termination codons (PTCs) have been published – all within the last exon and therefore predicted to escape nonsense-mediated decay (NMD) and result in a protein product (16). More recently, 4 nonsense and 3 frameshift variants that would be expected to result in NMD were uploaded to ClinVar (52), but it remains to be seen whether a reduction in EZH2 protein leads to the same phenotype as Weaver missense variants.

Since heterozygous whole-gene deletions can be assumed to reduce expression of the corresponding protein, we scanned for deletions encompassing *EZH2* in DECIPHER, a database linking human genetic variation with phenotype. Interestingly, of the 78 *EZH2* heterozygous whole-gene deletion variants with corresponding phenotype documentation available in DECIPHER, none are associated with overgrowth or tall stature (53). Rather, the second most common phenotype annotation is short stature, seen in 29 individuals (the top phenotype being intellectual disability, in 39 individuals). As these deletions range from 5.10 Mb to 27.36 Mb in length and span multiple genes, phenotypic effects cannot be attributed solely to the loss of *EZH2*. Indeed, a patient bearing a relatively smaller 1.2 Mb deletion has been reported to exhibit features of Weaver syndrome, such as tall stature, intellectual disability, and select facial dysmorphisms, which supports a mechanism of haploinsufficiency (54). However, the possibility remains that EZH2 haploinsufficiency in humans could constitute a distinct phenotype from missense variants. Without clinical suspicion for Weaver syndrome, such variants may escape detection, leading to underreporting.

### Both direct and indirect transcriptional effects may drive MDEM phenotypes

EM genes are exquisitely dosage-sensitive (55), and Ezh2 is the predominant mammalian methyltransferase for the transcriptionally silencing H3K27me3 mark. Therefore, we originally hypothesized that decreased Ezh2 activity would lead to widespread gene de-repression. However, there is no skew towards upregulation of gene expression in *Ezh2^R684C/+^* cells, suggesting that de-repressed Ezh2 target genes may be drowned out by innumerable downstream indirect effects. Despite the statistical significance of these gene expression changes, many of the corresponding fold-changes in expression were also relatively modest. In agreement with prior literature on MDEMs (6), we view these findings in light of Boyle et al.’s “omnigenic” model, which supposes that genes are highly interconnected regulatory networks (56). Perturbation of a few core genes, which have a direct biological link to disease, leads to small regulatory changes in a large number of interconnected genes. This is also concordant with our observation that of all the genes with altered expression, the great majority were not directly implicated in the osteogenic phenotype. The transcriptional profile of MDEMs such as Weaver syndrome can thus be thought of as the sum of direct effects due to locus-specific epigenetic modification as well as consequent indirect effects upon the gene regulatory network. Although outside the scope of this paper, concurrent ChIP for Ezh2 could distinguish direct gene targets from indirect; these may also be approximated by H3K27me3 ChIP-seq. This finer resolution may be required to parse out differences in H3K27me3 at individual loci, as we observed earlier that global H3K27me3 is minimally altered in heterozygous cells.

### Abnormal cell differentiation is a shared feature of MDEMs

Cell fate decisions are highly regulated processes in multicellular organisms. Epigenetic control of cell type-specific programs is critical to achieve spatiotemporally appropriate differentiation, and loss of this control has disastrous consequences (57). It is therefore unsurprising that the epigenomic disruption caused by MDEMs impacts cell differentiation. Our results in *Ezh2^R684C/+^* mice suggest that in Weaver syndrome, osteoblast differentiation is perturbed, at least in part through dysregulation of the BMP pathway. Excessive osteogenesis is concordant with the human phenotype of skeletal overgrowth and advanced osseous maturation. Studies of other MDEMs also indicate aberrant differentiation in phenotype-relevant tissues: precocious differentiation of chondrocytes and neural stem and progenitor cells (NSPCs) in Kabuki syndrome 1 (4, 5), and delayed maturation of NSPCs in Rubinstein-Taybi syndrome (MIM 180849) (58). Further investigation of other relevant cell types in Weaver syndrome may yield additional findings of altered differentiation.

### Epigenetic modifying agents such as GSK-J4 are a promising approach to treating MDEMs

No targeted therapies are yet approved for use in MDEMs, but advances are being made. Currently, MDEMs including Weaver syndrome are managed in clinic on the basis of ameliorating symptoms, which are often multi-systemic (3). Here, we sought a treatment to address the mechanistic root of Weaver syndrome. Fahrner and Björnsson previously proposed the ‘Balance hypothesis’, that opposing writers and erasers of histone marks exist in a balance to maintain the normal chromatin state (1). MDEMs are thought to disrupt this balance by inappropriately increasing or reducing levels of epigenetic modifications at target loci, leading to an abnormal chromatin state. Several groups have adopted a strategy of using epigenetic modifying agents to restore balance in MDEMs, specifically Rubinstein-Taybi syndrome and Kabuki syndrome 1 (59–62). Alarcón et al. first demonstrated that cognitive defects in a mouse model of Rubenstein-Taybi could be ameliorated by directly counteracting haploinsufficiency of the histone acetyltransferase, CBP, with histone deacetylase inhibitor (HDACi) (59). In the case of Kabuki syndrome 1, which is predicted to have loss of the transcriptionally activating mark H3K4me3 at promoters, the Björnsson group showed that either inhibition of the opposing H3K4 demethylase (62), or HDAC inhibition through pharmacological or diet-induced means was also sufficient to improve hippocampal function and visual-spatial learning in mice (60, 62). Therefore, correction of overall chromatin state may prove just as impactful as restoration of a specific epigenetic mark, albeit at a risk of lower specificity.

While the above studies focus on treatment of neurological features, our findings suggest a similar approach also has the potential to reverse skeletal overgrowth, another common feature of MDEMs. Here, we tested the ability of GSK-J4, an inhibitor of lysine demethylases Kdm6a and Kdm6b, to counteract the loss of Ezh2 function. The balance between the primary writer and erasers of H3K27me3 has previously been implicated in human bone biology (63, 64). Inhibition of Ezh2 by GSK-126, a pharmacological analogy to Weaver syndrome, stimulates osteoblast differentiation in mice (65), while GSK-J4 reduced osteogenesis in a mouse model of Saethre-Chotzen craniosynostosis (MIM 101400) (66). We showed that GSK-J4 not only reduces mineralization by *Ezh2^R684C/+^* BM-MSCs towards wild-type, but also substantially reverses effects of the variant allele at the transcriptional level. Of note, we previously found that correcting the expression of a single target gene was insufficient to restore normal chondrogenesis in Kabuki syndrome 1 (5). Here we showed that the transcriptional impact of GSK-J4 involved a fold-change reversal of over 1,000 genes, suggesting that a large network of pathogenic gene expression may need to be addressed in order to achieve phenotypic rescue in MDEMs. This finding stresses the importance of developing epigenetic modifying agents to treat MDEMs, as these drugs hold the prospect of enacting the broad transcriptomic changes required across a multitude of tissues. Our results constitute the first demonstration of using an epigenetic modifying agent to treat growth abnormalities in MDEMs, and strongly support the potential of GSK-J4 as a therapeutic for Weaver syndrome and possibly other overgrowth disorders impacting PRC2 function.

## Conclusion

Using our novel *Ezh2^R684C/+^* mouse model, we showed that osteoblasts are a key cell type contributing towards skeletal overgrowth in Weaver syndrome. We reversed the pathological osteogenic phenotype of *Ezh2^R684C/+^* osteoblasts *in vitro* with GSK-J4, an epigenetic modifying agent targeting Kdm6a/Kdm6b, the demethylases complementary to Ezh2. Future studies on the locus-specific role of H3K27 methylation in growth regulation are called for to shed light on molecular mechanisms of Weaver syndrome and related PRC2 syndromes. The ultimate goal of this work is to contribute towards the development of therapeutics to target the underlying etiology of MDEMs as a class of disorders.

## Methods

### Animals

*Ezh2^R684C/+^* mice were generated by the Johns Hopkins Transgenic Core Laboratory using CRISPR-Cas9 gene editing. The *Ezh2*-specific guide RNA (5’-GTGGTGGATGCAACCCGAAA-3’) and the homology-directed repair (HDR) template (5’-ACTGAAAATAAGTCACTGGATTATCTATGTTTTTCACTTTAGATTTTGTGGTGG ATGCAACatGtAAaGGCAACAAAATTCGTTTTGCTAATCATTCAGTAAATCCAAACTGCTATGCAAAA GGTA-3’) were based on mouse transcript NM_007971.2/ ENSMUST00000081721.12. The HDR template included mutations (indicated in lowercase) necessary to generate the mouse p.R679C missense change, which corresponds to human p.R684C. In addition, silent base changes were introduced to engineer an Nsp1 restriction site for rapid genotyping. C57BL/6J embryos were injected with Cas9 mRNA, the gRNA, and the HDR DNA oligo in the presence of SCR7. One founder was obtained and crossed to C57BL/6J, demonstrating germline transmission. Mice were backcrossed for 9 generations prior to experimental use in order to remove potential off-target sequence changes and maintained on a C57BL/6J background (Strain No. 000664, The Jackson Laboratory). Genotyping was performed by PCR and Sanger sequencing, in some cases accompanied by Nsp1 digest. Mice were cared for by Johns Hopkins Research Animal Resources. Mice were housed up to 5 per barrier cage in ventilated racks with HEPA-filtered and humidified air, under standardized light/dark cycles. Mice had ad libitum access to autoclaved feed (Envigo Teklad 2018SX) and reverse osmosis-filtered, hyperchlorinated water. Each cage was provided with autoclaved corncob bedding (Envigo Teklad 7092/7097) and a cotton square nestlet (Envigo Teklad 6060/6105). Cages were changed every 2 weeks under aseptic conditions. Euthanasia was performed by halothane inhalation (Sigma-Aldrich B4388), following the AVMA Guidelines for the Euthanasia of Animals, 2020 edition (https://olaw.nih.gov/policies-laws/avma-guidelines-2020.htm).

### Mouse embryonic fibroblasts (MEFs)

*Ezh2^R684C/+^* female mice underwent timed matings with *Ezh2^R684C/+^* male mice. Pregnant females were euthanized at E14.5. Embryos were dissected from the uteri, minced in DMEM with 15% FBS, L-glutamine, non-essential amino acids, and penicillin/streptomycin and dissociated with gentle pipetting. Cell suspension was cultured in DMEM with 10% FBS, L-glutamine, non-essential amino acids, and penicillin/streptomycin at 37°C, 5% CO_2_, with media changes every 48-72 hours.

### Mouse bone marrow mesenchymal stem cells (BM-MSCs)

Mice were euthanized at 8 or 10 weeks of age (males or females, respectively). Femora and tibiae were dissected immediately and rinsed with penicillin/streptomycin (ThermoFisher Scientific 15140122) in phosphate-buffered saline. Bone marrow was flushed out of the medullary cavity with BM-MSC complete growth media, consisting of MEM Alpha (Corning 10-022-CV) supplemented with 15% heat-inactivated fetal bovine serum (ThermoFisher Scientific 16140071), 2 mM L-glutamine (Corning 25-005-CI), 100 U/mL penicillin and 100 µg/mL streptomycin (ThermoFisher Scientific 15140122). Bone marrow was passed through a 70 µm cell strainer and cultured for 2 weeks in BM-MSC complete growth media at 37°C and 5% CO_2_. Media was refreshed every 48-72 hours. Adherent BM-MSCs were dissociated with 0.25% trypsin, 2.21 mM EDTA (Corning 25-053-CL) and seeded for subsequent experiments.

### Osteogenic differentiation

BM-MSCs were seeded at a density of 1×10^5^ cells/cm^2^ in BM-MSC complete growth media. After 24 hours, this was swapped for osteogenic differentiation media. Osteogenic differentiation media consisted of BM-MSC complete growth media supplemented with 0.05 mg/mL L-ascorbic acid (Sigma-Aldrich A4403), 10 mM β-glycerophosphoric acid (ThermoFisher Scientific 410991000), and 102 nM dexamethasone (Sigma-Aldrich D4902). Cells were continually cultured for 21 days at 37°C and 5% CO_2_. Osteogenic differentiation media was refreshed every 48-72 hours.

### Western blot

MEFs were washed twice with cold 1x PBS and lysed using RIPA buffer with a protease/phosphatase inhibitor cocktail (Cell Signaling Technology 5872) to obtain total protein samples. Histones were isolated with a Histone Extraction kit (Abcam ab113476). Protein concentration was assessed with the Pierce BCA Protein Assay kit (ThermoFisher Scientific 23225), and 20 µg of each sample was loaded onto pre-cast NuPAGE 4 to 12%, Bis-Tris gels (ThermoFisher Scientific NP0336BOX). Proteins were transferred to a PVDF membrane. 0.5x Intercept (PBS) Blocking Buffer (LI-COR Biosciences 927-70001) was used to block the membrane for 1 hour at room temperature. Membranes were incubated with the primary antibody overnight at 4°C, then washed using 1x PBS with 0.1% Tween-20 (PBS-T). Secondary antibody was added for 1 hour at room temperature. PBS-T was used to wash off unbound secondary. Membranes were imaged with the LI-COR Odyssey. All western blot antibodies are listed in Supplemental Table 2A.

### High-resolution micro-computed tomography

Femurs and tibias were dissected from 8-week old mice and fixed with 4% paraformaldehyde in 1x PBS for 48-72 hours before transfer to 70% ethanol. Bone length was assessed using digital calipers before high-resolution images were obtained with a Bruker Skyscan 1275 desktop microcomputed tomography system. Long bones were scanned at 65 keV and 152 µA using a 0.5 mm aluminum filter at an isotropic voxel size of 10 µm, in accordance with the recommendations of the American Society for Bone and Mineral Research (67). Micro-CT images were reconstructed with nRecon (Bruker) and analyzed with CTAn 3D analysis software (Bruker). Cortical bone structure was assessed in a 500 µm region of interest (ROI) centered on the femoral mid-diaphysis. Trabecular bone structure was assessed in a 2 mm ROI located 500 µm proximal to the distal femoral growth plate.

### Dynamic bone histomorphometry

5-week old mice were injected intraperitoneally with 10 mg/kg of calcein (Sigma-Aldrich C0875) dissolved in a 2% sodium bicarbonate solution, 5 days prior to euthanasia. Mice were then injected with 30 mg/kg of Alizarin red S (Sigma-Aldrich A3882) dissolved in a 2% sodium bicarbonate solution, 2 days prior to euthanasia. After collection, femurs were fixed in 100% ethanol until embedding in methyl methacrylate. Sections of 30 µm were obtained from the femoral mid-diaphysis and visualized with fluorescence microscopy. Histological analyses were performed using ImageJ in accordance with the American Society for Bone and Mineral Research (ASBMR) guidelines. Images were analyzed for endosteal (Es) and periosteal (Ps) mineral apposition rate (MAR), as defined by the ASBMR Committee for Histomorphometry Nomenclature (68, 69).

### Alizarin red staining

After 21 days of osteogenic differentiation, cells were washed with 1x PBS and fixed with 10% neutral buffered formalin (Epredia 9990244) for up to 1 hour. Following fixation, cells were washed with deionized water and stained with 2% Alizarin Red S solution, pH 4.2 (Electron Microscopy Sciences 26206-01) for 1 hour in the dark. Unbound dye was removed by repeated washes with deionized water. Whole-well images were taken over a transilluminator without magnification. The following protocol for quantification of Alizarin red was adapted from Serguienko et al., 2018 (70). Dye bound by the calcified matrix was dissolved by incubation in 10% acetic acid for 30 minutes with gentle agitation. Cells were scraped and transferred with the acetic acid solution to polypropylene tubes. Samples were heated to 85°C with agitation for 10 minutes, rapid-cooled on ice, then pelleted at 20,000 x g. Supernatant was retained and neutralized with 10% ammonium hydroxide to pH 4.1-4.5. Absorbance was measured at 405 nm with a BioTek Synergy 2 Multi-Mode Microplate Reader.

### MTT assay

CellTiter 96 AQueous One Solution (Promega G3580) was added to fresh BM-MSC complete growth media. Cells were incubated in the dark at 37°C, 5% CO_2_ for 1.5 hours to allow for conversion to formazan. Absorbance was measured at 490 nm with a BioTek Synergy 2 Multi-Mode Microplate Reader.

### qPCR

Total RNA was isolated from using TRIzol reagent (ThermoFisher Scientific 15596026). The SuperScript IV First-Strand Synthesis system (ThermoFisher Scientific 18091050) was used to reverse-transcribe the total RNA into cDNA. qPCR was performed with the PowerUp SYBR Green Master Mix (ThermoFisher Scientific A25742) on an Applied Biosystems ViiA 7 Real-Time PCR System. Primers are listed in Supplemental Table 2B.

### GSK-J4 treatment

GSK-J4 crystalline solid (Cayman 12073) was reconstituted in dimethyl sulfoxide (Sigma-Aldrich D2650) to a stock concentration of 10 mM. BM-MSCs were seeded at a density of 1×10^5^ cells/cm^2^ in BM-MSC complete growth media. After 24 hours, this was replaced by osteogenic differentiation media containing either GSK-J4 (treatment group) or the equivalent volume of DMSO (vehicle group). DMSO content did not exceed 0.1% of media volume for the highest treatment dose concentration. Cells were treated for 7 days, after which culture was maintained in osteogenic differentiation media without GSK-J4 or DMSO until day 21. Media was exchanged every 48-72 hours.

### RNA-sequencing library preparation

BM-MSCs were isolated from 6 mice each for *Ezh2^R684C/+^* and *Ezh2^+/+^* genotypes. BM-MSCs underwent osteogenic differentiation for 14 or 21 days. For the latter experiment, cells were treated with 2 µM of GSK-J4 or equivalent volume of DMSO (see Methods, GSK-J4 treatment). Cells were lysed and homogenized with TRIzol reagent (ThermoFisher Scientific 15596026), followed by phenol-chloroform separation. Total RNA was purified from the aqueous phase using the RNA Clean & Concentrator-5 kit with DNase I treatment to remove genomic DNA contamination (Zymo Research R1013). RNA quantity was determined with the Qubit RNA Broad Range Assay (ThermoFisher Scientific Q10210). RNA quality was assessed with the Agilent RNA 6000 Nano Kit (Agilent 5067-1511) run on an Agilent 2100 Bioanalyzer instrument, or submitted to the Johns Hopkins Single Cell & Transcriptomics Core (JH SCTC) Facility to be processed on an Agilent Fragment Analyzer. Polyadenylated RNA was isolated from either 1 µg (day 14) or 300 ng (day 21) of total RNA using the NEBNext Poly(A) mRNA Magnetic Isolation Module (New England BioLabs E7490L). Libraries were prepared using the NEBNext Ultra II RNA Library Prep kit with Sample Purification Beads (E7775S) and indexed with NEBNext Multiplex Oligos for Illumina (Dual Index Primers Set 1) (New England BioLabs E7600S). Library quality was assessed with the Agilent High Sensitivity DNA Kit (Agilent 5067-4626) run on the Agilent 2100 Bioanalyzer instrument or submitted to the JH SCTC for processing on an Agilent Fragment Analyzer. Completed libraries were quantified using the NEBNext Library Quant Kit for Illumina (New England BioLabs E7630L) and pooled accordingly to a final concentration of 4 nM. High-throughput sequencing was performed by the Johns Hopkins Genomics Research Core Facility on the Illumina NovaSeq 6000 platform using SP flow cells to generate 100-bp paired-end reads.

### Analysis of RNA-sequencing data

Salmon 1.9.0 was used to index the GRCm38 transcriptome (71), using the GRCm38 genome as the decoy sequence; both are available through Ensembl (Mus_musculus.GRCm38.cdna.all.fa.gz and Mus_musculus.GRCm38.dna.primary_assembly.fa.gz, from http://nov2020.archive.ensembl.org/Mus_musculus/Info/Index, release 102). Lane read outputs were demultiplexed into raw FASTQ files for each sample, which were mapped using Salmon 1.9.0 for paired-end reads, with selective alignment and GC bias correction enabled. Transcript quantifications were imported into R 4.1.2, running Bioconductor 3.14, and summarized to gene-level counts using tximeta 1.12.4 (72). Gene counts for any technical replicates were combined, as principal component analyses indicated minimal difference between technical replicates. Non- and low-expressed genes (median count across all samples < 10) were filtered. Surrogate variables (SVs) were identified and accounted for using sva 3.42.0 (2 SVs for D14; 4 SVs for D21) (73). DESeq2 1.34.0 was used to perform differential expression analysis (74). A false-discovery rate cutoff < 0.1 was used to determine significance; the default DESeq2 correction for multiple comparisons is the Benjamini-Hochberg procedure. Genes were subsequently annotated with biomaRt 2.50.3, accessing version 102 of the *Mus musculus* Ensembl dataset (75, 76). Lists of differentially expressed genes are available (Supplemental Appendices 1-3). The Mouse Genome Informatics database was accessed on March 25, 2023 to download lists of genes with the Gene Ontology annotations ‘BMP signaling pathway’ and ‘osteoblast differentiation’ (Supplemental Appendices 4 and 5). Two-sample Wilcoxon test statistics were calculated for the p-values of genes falling within and outside of the annotated subsets. These were compared to the Wilcoxon test statistic distribution calculated for 10,000 gene subsets of equal length, chosen at random.

### Validation of RNA-sequencing data using an external dataset

Raw read FASTQ files were downloaded from GEO accession GSE138980 (34). Samples treated with the EZH2 inhibitor GSK-126 were compared to samples treated with non-targeting siRNA (control group). Read mapping and differential expression analysis was performed as described in the ‘Analysis of RNA-sequencing data’ section of the Methods, maintaining a false-discovery rate cutoff < 0.1. A conditional p-value histogram was generated to examine the relative enrichment of differentially expressed genes (DEGs) in *Ezh2^R684C/+^* early osteoblasts among the pool of DEGs resulting from pharmacological inhibition of Ezh2 in murine mesenchymal stem cells.

### Determination of EZH2 target genes

The ‘Transcription Factor ChIP-seq Clusters (161 factors) from ENCODE with Factorbook Motifs’ track data was downloaded from the UCSC Genome Browser (wgEncodeRegTfbsClusteredWithCellsV3.bed.gz, http://hgdownload.soe.ucsc.edu/goldenPath/hg19/encodeDCC/wgEncodeRegTfbsClustered/) (77–81). EZH2-bound clusters were selected, and converted to a GRanges object using GenomicRanges 1.46.1 (82). The locations of all hg19 gene promoters (+/− 2 kb of the transcription start site) were determined using the reference genome package BSgenome.Hsapiens.UCSC.hg19 alongside the annotation package EnsDb.Hsapiens.v75. A gene was considered an EZH2 target if its promoter overlapped with an EZH2-bound cluster. biomaRt 2.50.3 was used to match hg19 EZH2 target genes with mouse homologs (75, 76).

### Statistics

Two-tailed, unpaired Student’s t-tests were used for most analyses (excepting RNA-seq studies), with p < 0.05 as the threshold for statistical significance. Where applicable, the Benjamini-Hochberg procedure was used to correct for multiple comparisons. Statistics for the RNA-seq analyses are detailed under the Methods section ‘Analysis of RNA-sequencing data’.

### Study approval

All animal procedures and protocols in this study were approved by the Johns Hopkins Institutional Animal Care and Use Committee, and performed in accordance with the NIH *Guide for the Care and Use of Laboratory Animals* (National Academies Press, 2011).

### Data availability

[The data discussed in this publication are in the process of being deposited in NCBI’s Gene Expression Omnibus (83) and accession number will be made available, once received. Analysis code will be made available on GitHub.]

## Author contributions

JAF designed the CRISPR-Cas9 edit to generate the *Ezh2^R684C/+^* mouse and isolated MEFs. WYL and JAF performed the western blot and initial mouse phenotyping. PK and RR collected and analyzed the micro-CT and dynamic bone histomorphometry data. WYL optimized conditions for osteoblast differentiation and prepared the libraries for untreated RNA-seq. CWG performed GSK-J4 treatments and library preparation for drug-treated RNA-seq. CWG, LB, and KDH analyzed the RNA-seq data. CWG wrote the manuscript. All authors contributed towards the editing and approval of this manuscript. HTB contributed essential resources at the start of the project. HTB and KDH participated in helpful discussions throughout. JAF conceived and directed the project and oversaw its completion.

## Acknowledgments

We thank the following: Roger Reeves, Chip Hawkins, Holly Wellington, and Ann Lawler in the Johns Hopkins Transgenic Mouse Core Facility for their assistance generating the *Ezh2^R684C/+^* mouse model; Tom Clemens, Vinod Ranganathan, Joel Pomerantz, and Corinne Hamblet for helpful discussions; Li Zhang for technical assistance; Jake Volk in the Johns Hopkins Single Cell and Transcriptomics Core Facility; David Mohr at the Johns Hopkins Genetic Resources Core Facility; Kim Sealover for help with mouse husbandry; Bailey Spiegelberg for researching the Bmp pathway. We thank Carol W Greider for providing helpful feedback on the manuscript. We acknowledge the ENCODE Consortium for use of ENCODE data. The Factorbook Motifs track data was generated by the labs of Richard Myers, Michael Snyder, Mark Gerstein, Sherman Weissman, Peggy Farnham, Kevin Struhl, Kevin White, and Vishy Iyer. This study also makes use of data generated by the DECIPHER community. A full list of centers who contributed to the generation of the data is available from https://deciphergenomics.org/about/stats and via email from. Funding for the DECIPHER project was provided by Wellcome grant number WT223718/Z/21/Z. The research in this manuscript was primarily funded by K08HD086250, awarded by the National Institutes of Health (NIH)/National Institute for Child Health and Human Development (NICHD; JAF), a Hartwell Foundation Individual Biomedical Research Award (JAF), a Johns Hopkins Clinician-Scientist Award (JAF), a grant from the William and Ella Owens Medical Research Foundation (JAF), and Merit Review Grant BX003724 from the Biomedical Laboratory Research and Development Service of the Veterans Affairs Office of Research and Development (RCR). HTB is supported by funding from The Icelandic Centre for Research (#217988, #195835, #206806) and the Louma G. Foundation. CWG is supported by NIGMS T32GM136577 and a JHU Bloomberg Distinguished Professorship to Carol W Greider.

## Notes

### Competing Interest Statement

HTB is a consultant for Mahzi therapeutics. The other authors have no conflicts of interest.

### Summary of Updates

Figure 1C corrected.

