## Supplemental Figures and Tables for "Novel mouse model of Weaver syndrome displays overgrowth and excess osteogenesis reversible with KDM6A/6B inhibition"

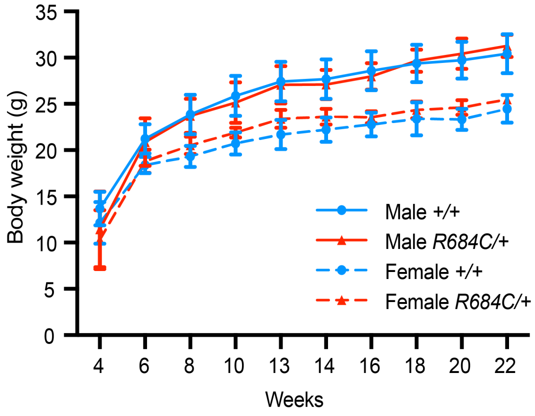


**Supplemental Figure 1. Growth curves of *Ezh2*$^{R684C/+}$ mice and *Ezh2*$^{+/+}$ littermates from 4-22 weeks of age.** Female *Ezh2*$^{R684C/+}$ mice trend towards a higher weight than *Ezh2*$^{+/+}$ littermates from 8 weeks of age onward. Blue circles with solid lines represent male *Ezh2*$^{+/+}$ (n=8); red triangles with solid lines represent male *Ezh2*$^{R684C/+}$ (n=5); blue circles with dashed lines represent female *Ezh2*$^{+/+}$ (n=5); red triangles with dashed lines represent female *Ezh2*$^{R684C/+}$ (n=4). All error bars represent mean ±1 SD.

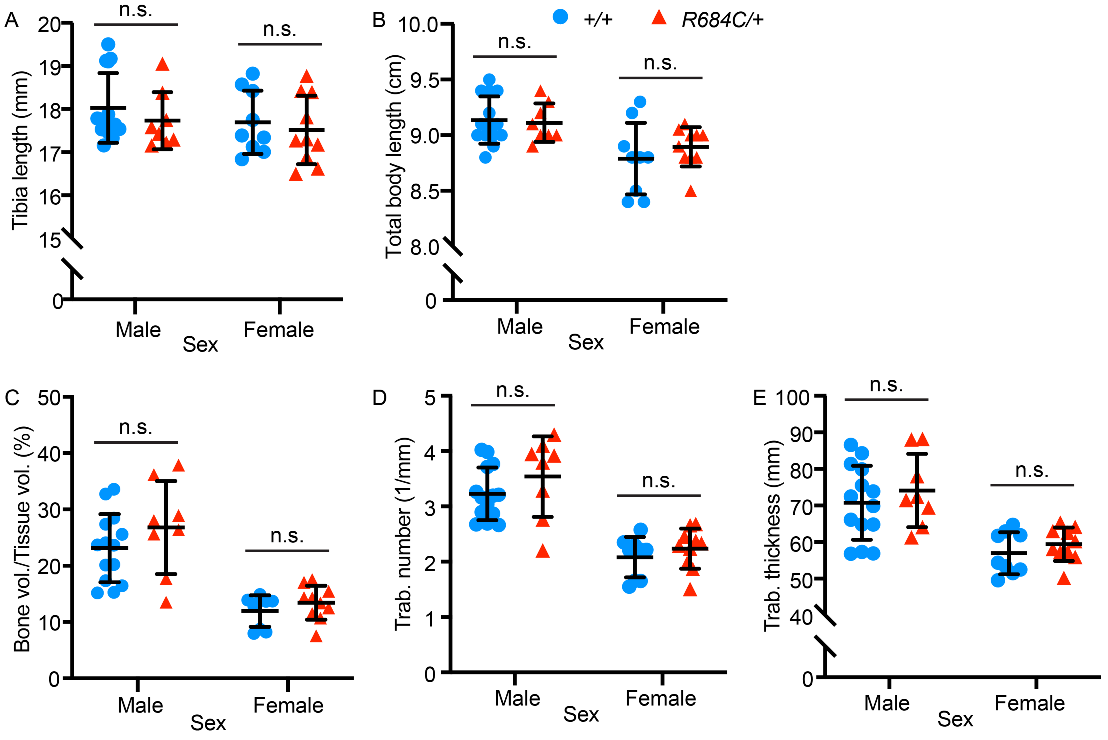


**Supplemental Figure 2. Additional bone measurements and micro-CT parameters.** (A) Tibia length and (B) total body length do not differ between *Ezh2*$^{R684C/+}$ and *Ezh2*$^{+/+}$ mice for either sex. (C) Trabecular bone volume (bone volume/tissue volume), (D) trabecular number, and (E) trabecular thickness do not differ between *Ezh2*$^{R684C/+}$ and *Ezh2*$^{+/+}$ femurs. Blue circles represent *Ezh2*$^{+/+}$. Red triangles represent *Ezh2*$^{R684C/+}$. All error bars represent mean ±1 SD. Trab., trabecular; n.s., non-significant.

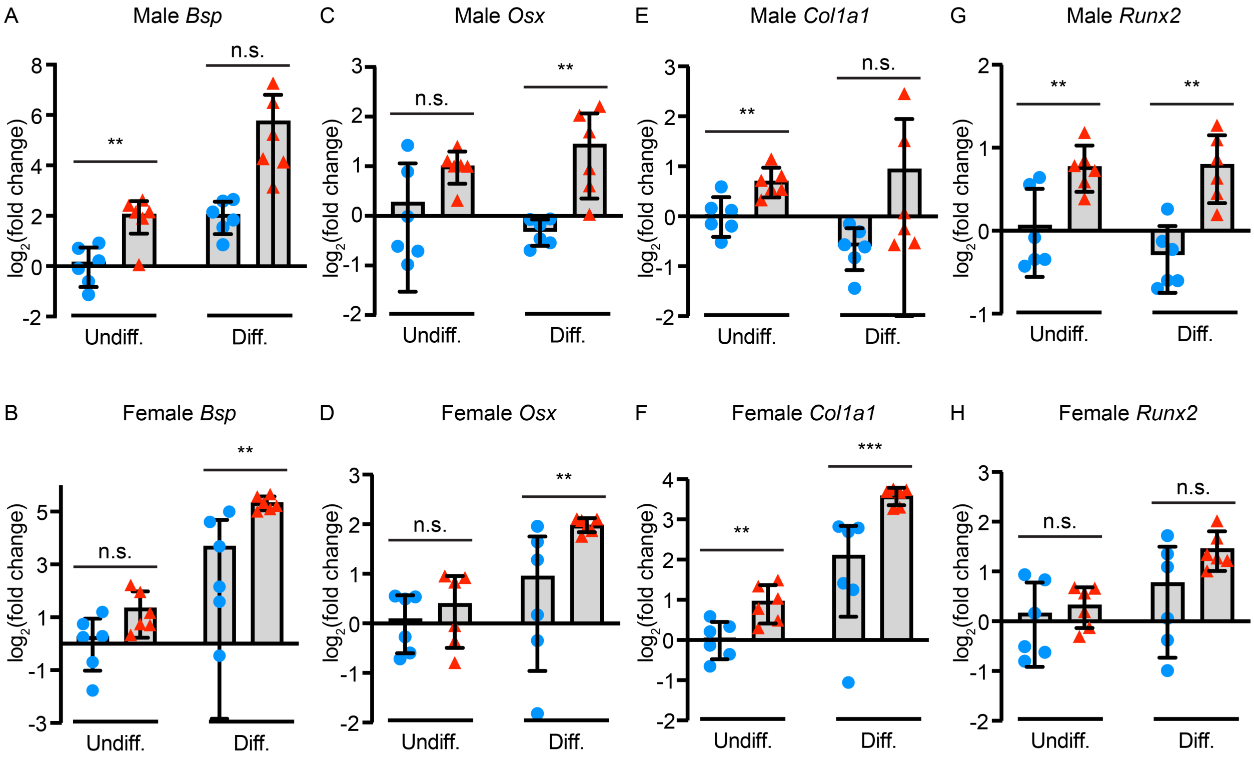


**Supplemental Figure 3. Markers of osteogenesis are upregulated in *Ezh2*$^{R684C/+}$ undifferentiated BM-MSCs and differentiated osteoblasts.** qPCR for *Bsp* in (A) males and (B) females; *Osx* in (C) males and (D) females; *Col1a1* in (E) males and (F) females; and *Runx2* in (G) males and (H) females. RNA was isolated from *Ezh2*$^{+/+}$ and *Ezh2*$^{R684C/+}$ cells (n=6 per genotype), in either an undifferentiated BM-MSC state, or after differentiation towards osteoblasts. For all plots, blue circles represent *Ezh2*$^{+/+}$; red triangles represent *Ezh2*$^{R684C/+}$. $^{**}p < 0.01$, $^{***}p < 0.001$, unpaired Student's t-test. All error bars represent mean ±1 SD. Undiff., undifferentiated. Diff., differentiated; n.s., non-significant.

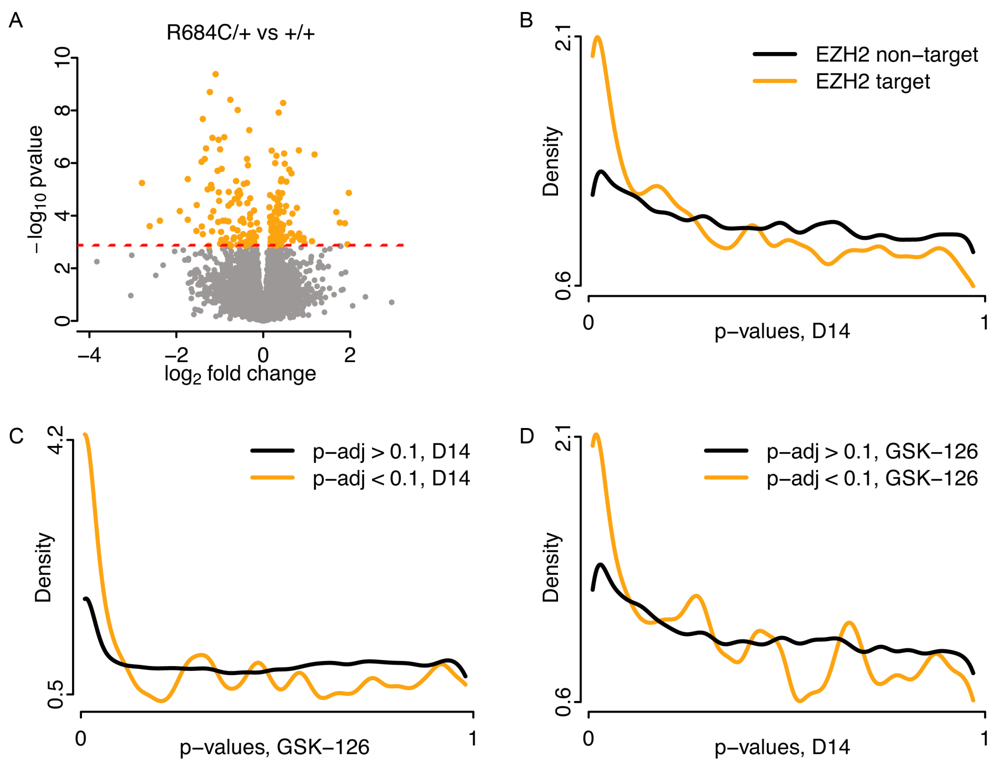


**Supplemental Figure 4. RNA-seq comparing *Ezh2*$^{R684C/+}$ osteoblasts to *Ezh2*$^{+/+}$ after 14 days of differentiation, and validation with publically available datasets.** (A) Volcano plot of RNA-seq performed on osteoblasts differentiated from *Ezh2*$^{R684C/+}$ and *Ezh2*$^{+/+}$ BM-MSCs for 14 days. Differentially expressed genes are highlighted in orange. False discovery rate (FDR) = 0.1 (red dashed line). The ENCODE project aggregated ChIP-seq data for various transcription factors, including EZH2, in various human cell lines. Because EZH2 is highly conserved between human and mouse, we assumed that Ezh2 would bind at orthologous target promoters in mouse. Furthermore, we reasoned that genes bound and regulated by Ezh2 would be most impacted by a reduction in Ezh2 function. This is the case, as shown in the (B) conditional p-value density plot of p-values comparing *Ezh2*$^{R684C/+}$ to *Ezh2*$^{+/+}$ at Day 14, stratified by whether or not the gene is a human target of EZH2 (orange line indicates EZH2 target; black line indicates EZH2 non-target). Sen et al. previously published an RNA-seq dataset generated from murine bone marrow mesenchymal stem cells (BM-MSCs). Prior to harvest, the cells underwent osteoblast differentiation for 3 days while being treated with a scramble siRNA (control) or the EZH2 inhibitor GSK-126. We reasoned that the transcriptional profile of BM-MSCs treated with GSK-126 should bear similarities with that of *Ezh2*$^{R684C/+}$ mice, as both have reduced Ezh2 functionality. (C) Conditional p-value density plot showing p-values from differential expression due to GSK-126 treatment, stratified by significance in our Day 14 RNA-seq which compared *Ezh2*$^{R684C/+}$ to *Ezh2*$^{+/+}$ (orange line indicates p-adj < 0.1; black line indicates p-adj > 0.1). The relative enrichment of low p-values for genes marked as significant in the Day 14 dataset indicates a degree of concordance in the genes most affected by Ezh2 dysfunction, whether induced genetically or pharmacologically. (D) Conditional p-value density plot of p-values from the Day 14 RNA-seq, stratified by significance in the GSK-126 dataset.

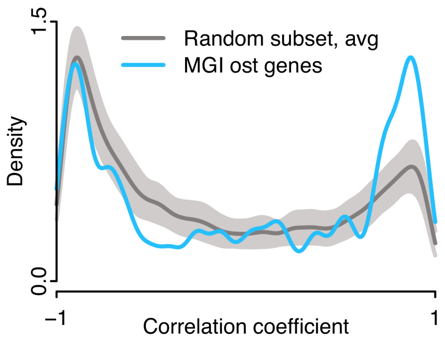


**Supplemental Figure 5. Osteoblast differentiation genes correlate strongly with PC1 of Day 14 RNA-seq comparing *Ezh2*$^{R684C/+}$ osteoblasts to *Ezh2*$^{+/+}$.** Density plot of correlation coefficients between sample PC1 coordinates and sample reads for each gene from the Day 14 RNA-seq data (normalized for library size as counts per million). Blue line represents density plot of PC1 correlation coefficients for the 179 MGI osteoblast differentiation genes. Dark gray line represents mean density plot of PC1 correlation coefficients for 179 genes selected at random, iterated 100 times. Light gray shading represents ±1 SD from the mean density for 100 iterations. Ost., osteoblast differentiation.

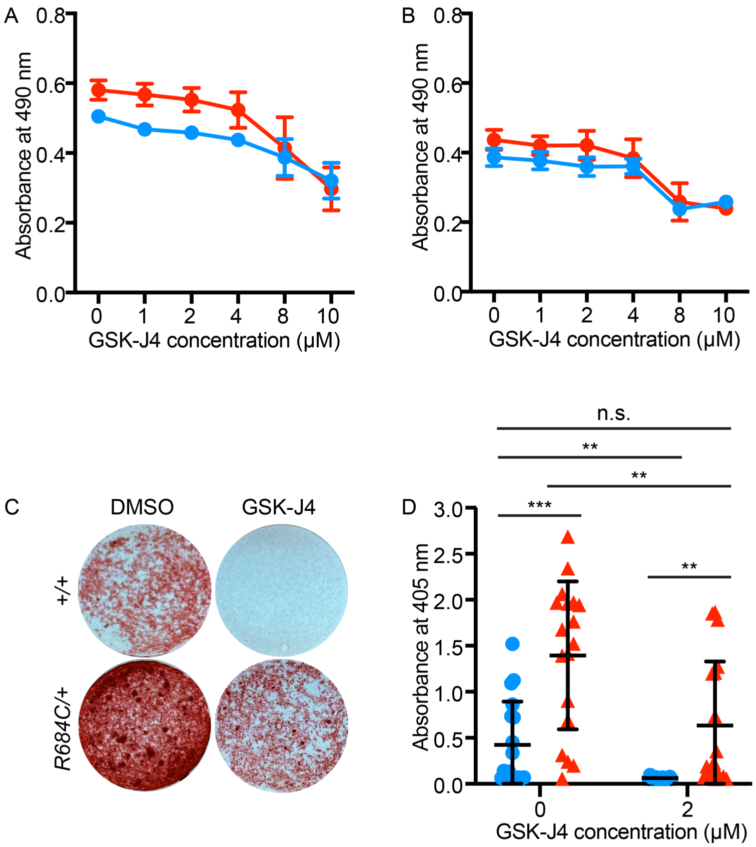


**Supplemental Figure 6. Determination of GSK-J4 dosage, and response to GSK-J4 treatment in male osteoblasts.** (A) MTT assay to measure cell viability after treatment with varying dosages of GSK-J4 for 7 days. Shown are mean absorbance values at 490 nm for BM-MSCs derived from 6 female *Ezh2*$^{+/+}$ mice (blue circles) and 6 female *Ezh2*$^{R684C/+}$ mice (red triangles). (B) This was also repeated with BM-MSCs isolated from male mice, n=6 per genotype. (C) Representative whole well images from Alizarin red staining of male *Ezh2*$^{+/+}$ and *Ezh2*$^{R684C/+}$ osteoblasts at Day 21. Cells were treated with either 2 μM of GSK-J4 or an equivalent volume of DMSO (vehicle) for the first 7 days. (D) Alizarin red quantification, showing that *Ezh2*$^{R684C/+}$ osteoblasts had higher absorbance at 405 nm than *Ezh2*$^{+/+}$ osteoblasts in either the DMSO or GSK-J4 condition. For both genotypes, GSK-J4 treatment led to a significant decrease in dye uptake, indicating lower osteogenic activity. Additionally, GSK-J4-treated *Ezh2*$^{R684C/+}$ osteoblasts had no significant difference from DMSO-treated *Ezh2*$^{+/+}$ osteoblasts. n=18 for each genotype. **$p < 0.01$, ***$p < 0.001$, unpaired Student's t-test with Benjamini-Hochberg procedure to correct for multiple comparisons. All error bars represent mean ±1 SD. n.s., non-significant.

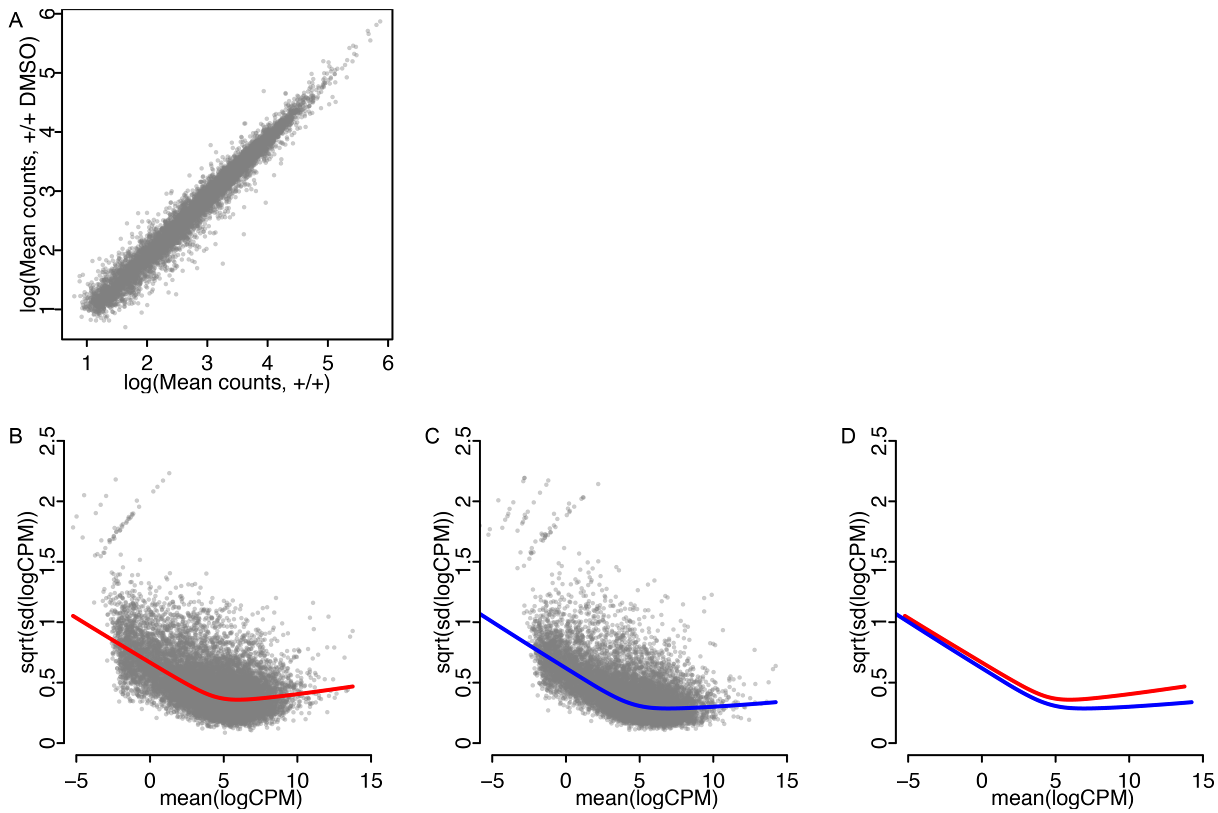


**Supplemental Figure 7. Differences between Day 14 and Day 21 RNA-seq results are not due to count irregularities or increased variance.** (A) Scatter plot comparing log(mean counts) from Day 14 RNA-seq for +/+ samples and log(mean counts) from Day 21 RNA-seq for +/+ DMSO samples. Each point represents one gene. Mean-variance relationships were plotted with LOWESS trend lines for (B) Day 14 +/+ samples and (C) Day 21 +/+ DMSO samples. (D) Overlay of LOWESS trend lines for Day 14 +/+ samples (red) and Day 21 +/+ DMSO samples (blue).

**Supplemental Table 1:**

A. *R684C/+* x *R684C/+* (at term)

| Cross | +/+ | *R684C/+* | *R684C/R684C* |
|---|---|---|---|
| 1 | 2 | 6 | 0 |
| 2 | 1 | 5 | 0 |
| 3 | 2 | 10 | 0 |
| 4 | 1 | 12 | 0 |
| **Total** | **6** | **33** | **0** |

B. *R684C/+* x *R684C/+* (E14.5)

| Cross | +/+ | *R684C/+* | *R684C/R684C* |
|---|---|---|---|
| 1 | 1 | 2 | 4 |
| 2 | 3 | 5 | 1 |
| **Total** | **4** | **7** | **5** |

C. *R684C/+* x +/+ (at term)

| Cross | +/+ | *R684C/+* |
|---|---|---|
| 1 | 18 | 12 |
| 2 | 9 | 10 |
| 3 | 14 | 6 |
| 4 | 8 | 13 |
| **Total** | **49** | **41** |

**Supplemental Table 2:**

A. Western antibodies:

| | Species | Manufacturer | Catalog number |
|---|---|---|---|
| anti-H3K27me3 | Rabbit | Millipore Sigma | 07-449 |
| anti-H3 | Rabbit | Cell Signaling Technology | 4499 |
| anti-Ezh2 | Rabbit | Cell Signaling Technology | 5246 |
| anti-β-actin | Mouse | Cell Signaling Technology | 3700 |
| IRDye 800CW Donkey anti-Rabbit IgG | Donkey | LI-COR | 926-32213 |
| IRDye 680RD Donkey anti-Mouse IgG | Donkey | LI-COR | 926-68072 |

B. qPCR primers:

| Target | Forward | Reverse |
|---|---|---|
| *Bsp* | 5'-ATGGAGACGGCGATAGTTCC-3' | 5'-CTAGCTGTTACACCCGAGAGT-3' |
| *Osx* | 5'-ATGGCGTCCTCTCTGCTTG-3' | 5'-TGAAAGGTCAGCGTATGGCTT-3' |
| *Col1a1* | 5'-GCTCCTCTTAGGGGCCACT-3' | 5'-CCACGTCTCACCATTGGGG-3 |
| *Runx2* | 5’-TTCAACGATCTGAGATTTGTGGG-3’ | 5’-GGATGAGGAATGCGCCCTA-3’ |
| *Hprt* | 5'-CTGGTGAAAAGGACCTCTCGAAG-3' | 5'-CCAGTTTCACTAATGACACAAACG-3' |
